# Adaptive evolution of *Plasmodium vivax* in Duffy-negative hosts: insights from East African genomes

**DOI:** 10.1101/2025.11.13.688259

**Authors:** Anthony Ford, Cheikh Cambel Dieng, Beka Raya Abagero, Guiyun Yan, Delenesaw Yewhalaw, Daniel A. Janies, Eugenia Lo

## Abstract

Historically, *Plasmodium vivax* (*Pv*) malaria was rare in Africa due to the lack of Duffy antigen receptor for chemokines (DARC) expression on host erythrocytes, an essential receptor for *Pv* Duffy Binding Protein (PvDBP1)-mediated invasion. However, increased reports of *Pv* cases across Africa and among Duffy-negative individuals have led to the hypothesis that the parasites have evolved alternate invasion mechanisms that are DARC-independent. To investigate potential genetic adaptations underlying this phenomenon, we performed genome-wide single nucleotide polymorphisms (SNPs) analyses of 110 *Pv* isolates collected from Ethiopia, comprising 38 Duffy-negative and 72 Duffy-positive infections. *Pv* from Duffy positives exhibited markedly higher genetic variation (477,561 SNPs) than those from Duffy negatives (197,461 SNPs). Chromosomes 1, 2, 9, 10, and 12 harbored the highest SNPs densities, consistent across both host genotypes but elevated in Duffy-positive infections. Among 43 erythrocyte-binding gene candidates, tryptophan rich antigen 3 and 34 (*TRAg*3 and *TRAg*34) and members of the merozoite surface protein 3 (*MSP*3) family showed the greatest nucleotide diversity per kilobase, highlighting these loci as potential mediators of host-parasite interaction shifts. Signals of positive selection differed by host Duffy genotypes. In Duffy-positive *Pv*, adaptive signatures were observed in genes related to drug resistance (chloroquine resistance-associated protein *CG*1 and 26S proteasome regulatory subunit *RPN*2) and erythrocyte binding (*MAEBL*); whereas in Duffy negative *Pv*, positive selection was observed in genes linked to organellar maintenance and vesicle trafficking (plastid replication repair enzyme and the AP-5 complex subunit beta 1), implicating alternative metabolic or trafficking adaptations. Amino acid substitutions in invasion ligands (*PvDBP*1, *PvEBP/DBP*2, and *PvRBP*2b) were common in Duffy-positive *Pv* but largely absent in Duffy-negative ones, with almost half of the mutations located in critical binding domains. Overall, *Pv* isolates in Duffy-negative hosts displayed reduced genomic diversity yet retained high conservation across the genome, suggesting strong selective constraints and limited diversification. Phylogenetic comparison revealed that Ethiopian *Pv* clustered closely with other East African isolates, whereas Southeast Asian and South American *Pv* represented more distant lineages. These findings indicate that *Pv* circulating in Duffy-negative populations maintains a genetically conserved background, potentially reflecting stringent evolutionary bottlenecks and/or specialized host-parasite interactions required for invasion of Duffy-negative erythrocytes.

## Introduction

*Plasmodium vivax* (*Pv*) and *P. falciparum* (*Pf)* are two major human malaria parasite species worldwide. While *Pf* is the most lethal, *Pv* poses a significant health burden due to its widespread distribution across Southeast Asia, South America, the Middle East, and increasingly Africa. In Africa, reports of *Pv* cases now span Ethiopia, Botswana, Cameroon, Angola, Equatorial Guinea, Senegal, Sudan, and beyond (1–4). Despite an estimated 2 billion people at risk, *Pv* malaria is often overlooked as a neglected tropical disease because of its relatively low mortality compared to *Pf* (5).

Historically, populations of African descent were thought to be protected from *Pv* due to the lack Duffy Antigen Receptor for Chemokines (DARC) expression, a receptor essential for *Pv* Duffy Binding Protein (PvDBP1)-mediated invasion (6). However, several recent studies confirmed *Pv* infections in Duffy-negative individuals broadly across Africa, including areas where Duffy-negatives predominate, challenging the long-standing paradigm of Duffy-dependent invasion. Although such infections may present with lower parasitemia or reduced clinical severity, the successful adaptation and transmission of *Pv* in Duffy-negative populations demand a reevaluation of malaria surveillance, treatment, and elimination strategies in Africa (7, 8).

The *Pv* genome exhibits extensive polymorphisms, reflecting the combined effects of gene flow due to human migration, transmission intensity, and host susceptibility (9). In *Pf,* several erythrocyte receptor-ligand pathways support erythrocyte invasion, but in *Pv,* invasion is primarily mediated through PvDBP1-DARC interactions. Our previous study showed that *PvDBP1* mutations unique to *Pv* in Duffy-negative Ethiopians did not lead to binding of Duffy-negative erythrocytes *in vitro* (7, 10). Yet, other ligands such as *PvRBP1a*, *PvRBP1b*, *PvRBP2a* and *PvRBP2*-P2 in the reticulocyte binding protein (*PvRBP*) gene family play pivotal roles in reticulocyte invasion, with *PvRBP2b* binding to transferrin receptor 1 (TfR1) prior to invasion (11–16). Likewise, the Long Homology Repeat A (LHR-A) subdomain of the complement receptor 1 (CR1) was recently shown to bind to region II of *PvEBP/DBP2* (17). Furthermore, highly polymorphic genes in the *PvMSP* and *PvTRAg* multigene families have been implicated in both erythrocyte invasion and immune evasion (18–21). Genetic diversification within these loci could therefore modulate invasion efficiency, host specificity, and adaptation to Duffy-negative erythrocytes. Understanding how invasion ligands evolved is critical for elucidating the molecular basis of DARC-independent invasion and its contribution to expanding range of *Pv* in Africa.

Recent population analysis of 730 global *Pv* isolates showed reduced genetic diversity and elevated linkage disequilibrium in the African isolates relative to those from Southeast Asia and South America, consistent with an Asian origin followed by sequential founder events during continental spread (22–24). Within Africa, *Pv* from Eritrea and Sudan clustered together and were distinct from the Ethiopian *Pv*, suggesting local adaptation (23). It is yet unclear the level of polymorphisms and the evolutionary pressures acting on invasion-related genes in *Pv* from Duffy negative Africans. Because of the highly polymorphic nature of *Pv*, genetic differences between Duffy-negative and Duffy-positive individuals may reflect a combination of adaptive mutations driven by local selection, transmission intensity, and host genetic background. In this study, we performed a genome-wide comparison of *Pv* from Duffy-positive and Duffy-negative hosts in Ethiopia, with the goals to characterize polymorphisms, determine variants across 43 erythrocyte-binding gene candidates, and infer evolutionary trajectories of African *Pv* lineages (9). Our findings illuminate the genomic underpinnings of alternate invasion mechanisms and provide new insight into the evolutionary origins and adaptive strategies of *Pv* in Africa.

## Materials and Methods

### Ethics Statement

Scientific and ethical clearance was obtained from the Institutional Scientific and Ethical Review Boards of Jimma University, Ethiopia and Drexel University, USA. Written informed consent/assent for study participation was obtained from all consenting heads of households, parents/guardians (for minors under 18 years old), and each individual who was willing to participate in the study.

### Sample Processing and Whole Genome Sequencing

A total of 110 *Pv*-confirmed whole blood samples were included, of which 38 were from Duffy-negative and 72 from Duffy-positive patients. These samples were monoclonal infections, as confirmed by microsatellite genotyping. We used a Lymphoprep^TM^/Plasmodpur^TM^-based protocol to deplete white blood cells and enrich red blood cell (RBC) pellets. Genomic DNA was extracted from approximately 1 mL of RBC pellets using Quick-DNA^TM^ Miniprep Kit (Zymo Research), following the manufacturer’s protocol. DNA quality and concentration were assessed using NanoDrop^TM^ spectrophotometer. DNA library was prepared using the NEBNext® Ultra^TM^ II DNA Library Prep Kit (New England BioLabs) following the manufacturer’s protocol for fragmentation, end repair, ligation, and PCR enrichment. Library quality and quantity were evaluated using Qubit fluorometry and Agilent TapeStation. Sequencing was carried out on the Illumina NovaSeq^TM^ 6000 platform, targeting a sequencing depth of 30X per sample to ensure comprehensive coverage of the genome. Quality control metrics included average Phred quality scores above 30 (Q30) across all bases and all samples that met the minimum threshold for both read depth and quality, ensuring reliable downstream analysis. Illumina sequencing reads were trimmed using Trimmomatic v0.39.

Whole genome sequencing (WGS) data of an additional 177 *Pv* samples were retrieved from other studies (25, 26), representing diverse geographical regions including Southeast Asia (Thailand, Cambodia), South America (Peru, Brazil, Guyana), and other parts of Africa (Uganda, Sudan, Madagascar, and Eritrea). All FASTQ files were mapped to the P01 reference genome, using BWA-mem v2 with default parameters (27–30). Compared to the Sal1 reference genome, P01 offers improved annotation especially in the subtelomeric regions of the *Pv* genome (29). Post-mapped reads were further processed to include sorting and flagging duplicate reads using Samtools v1.10 and Picard (31, 32). Only reads mapped to the reference genome were included in further analyses. All genome data were deposited in NCBI SRA database (PRJNA1255830).

### Variant Discovery, Nucleotide Diversity, and Codon Prevalence

Variant call format (VCF) files were generated using GATK’s HaplotypeCaller procedure with the best practices workflow (33). To ensure high-confidence variant calls, insertions and deletions were excluded from further analysis and SNPs were retained only if the following quality thresholds were met: QD (quality by depth) less than 2.0, QUAL (read quality) less than 30, SOR (strand odds ratio) greater than 3·0, and MQ (map quality) less than 40. Additionally, the latest version of gene annotation files was retrieved from PlasmoDB and functional annotations were applied to the remaining high-quality variants using snpEff v5.1 (30).

Nonsynonymous and synonymous variants were compared between the two Duffy groups across the 14 chromosomes in the nuclear genome. Coding sequences for 43 erythrocyte-binding genes, including members of the *PvTRAg*, *PvMSP*, *PvRBP*, *PvGAMA*, and *PvDBP* multigene families, were extracted from the assembled genomes (9). Nucleotide diversity was calculated for each gene and compared between Duffy-positive and Duffy-negative samples, excluding gaps from pairwise comparison. To control for differences in sample size between the two Duffy groups, we employed a bootstrap resampling approach to assess the robustness of nucleotide diversity calculations. For each gene, a fixed number of samples (*n*=38, the number of Duffy-negative samples) were randomly sampled with replacement from the Duffy-positive population. This procedure was repeated 500 times for each gene in each bootstrap replication to calculate nucleotide diversity. All resampling, computation and data aggregation were implemented in a customized Python script using pandas and NumPy. Amino acid variation for erythrocyte-binding genes including *PvTRAg38*, *PvDBP1*, *PvEBP/DBP2*, *PvMSP1*, *PvRBP1a*, *PvRBP1b*, and *PvRBP2a* was calculated using a customized python script, with the relative frequency of substitutions estimated for each codon. Substitutions within the binding domains were compared between Duffy-positive and Duffy-negative populations using Fisher’s exact test. To account for multiple testing across multiple sites, *p*-values were adjusted using the Benjamini-Hochberg correction.

### Test for signal of positive selection

We detected loci under recent positive selection using the integrated haplotype score (iHS), which compares the decay of extended haplotype homozygosity between reference and non-reference alleles at each SNP site, implemented in the R-based package *rehh* (*34, 35*). Because ancestral information was unavailable, the reference and non-reference alleles were used in place of the ancestral and derived alleles. We used a two-sided *p*-value for significance testing under the null hypothesis of no evidence of signals of recent selection. SNPs exhibiting iHS scores that had a *p*-value of −log_10_(*p*-value) greater than four were considered significant and interpreted as signature of positive selection.

### Population structure and differentiation

Population genetic analysis was assessed using the Admixture software, which estimates individual ancestry proportions using an accelerated maximum likelihood estimation by means of a novel quasi-Newton acceleration method (36). The analysis used a combination of the 110 Duffy-positive and Duffy-negative Ethiopian genomes alongside 177 publicly available genomes representing diverse global regions (36). Bayesian analyses was performed from 1 to 6 clusters (*K*), and cross-validation scores were calculated for each run, with the optimal *K* being the smallest cross-validation error.

### Statistical Analyses

Differences in SNP density at the chromosome and gene levels between Duffy-positive and Duffy-negative groups were assessed separately for nonsynonymous and synonymous mutations using the nonparametric Mann-Whitney U test. For each test, the null hypothesis was that the distributions of SNPs per 10 kb (chromosome level) and SNPs per kb (gene level) were the same between the two Duffy groups. All statistical tests were two-sided, and a *p*-value ≤ 0.05 was considered statistically significant. Statistical analyses were performed using Python with the scipy library.

## Results

### Genome-wide SNP variation reveals reduced diversity in Duffy-negative *Pv*

A total of 477,561 high quality SNPs were detected in *Pv* from Duffy-positive individuals, more than two-fold higher than the 197,461 SNPs observed in Duffy negatives. Among all observed SNPs, 134,266 were shared between the two Duffy groups, while 343,295 were unique to Duffy-positive and 63,195 were unique to Duffy-negative *Pv*. Across the genomes, the highest SNP density was observed in chromosomes 1, 2, 9, 10, and 12 in *Pv* of Duffy-positive individuals (Figure 1). Similar pattern was observed in the Duffy negatives, despite a lower number of nonsynonymous to synonymous SNPs. Though the number SNPs per 10 kb were higher in Duffy-positive than in Duffy-negative individuals, the Mann-Whitney U test showed no significant difference in the density of synonymous (*p*=0.533) and nonsynonymous (*p*=0.19) SNPs between Duffy-positive and Duffy-negative groups. These results suggest similar selective constraints across both Duffy groups, despite the overall reduction in SNP variants in Duffy-negative *Pv*.

**Figure 1.**
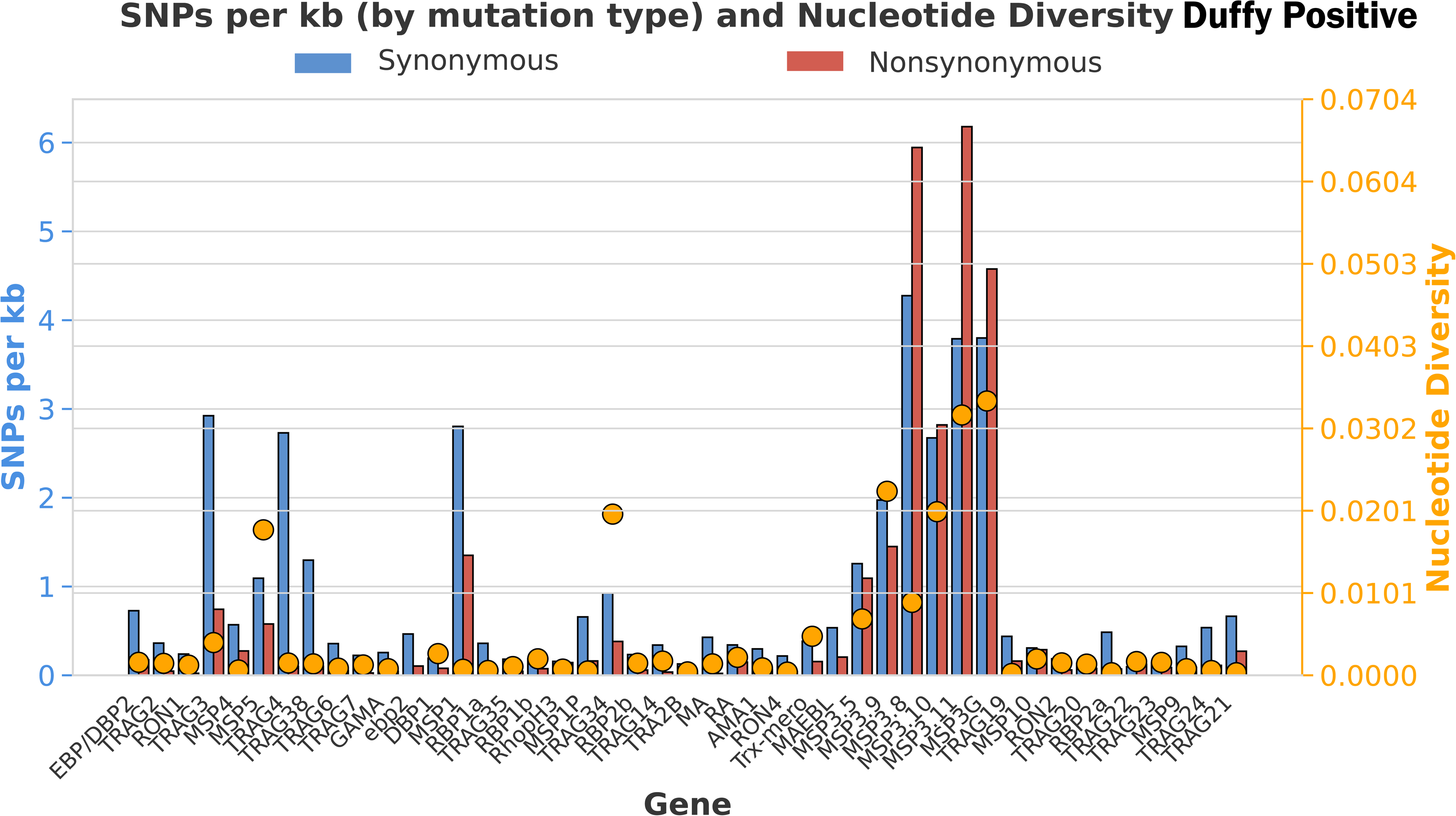
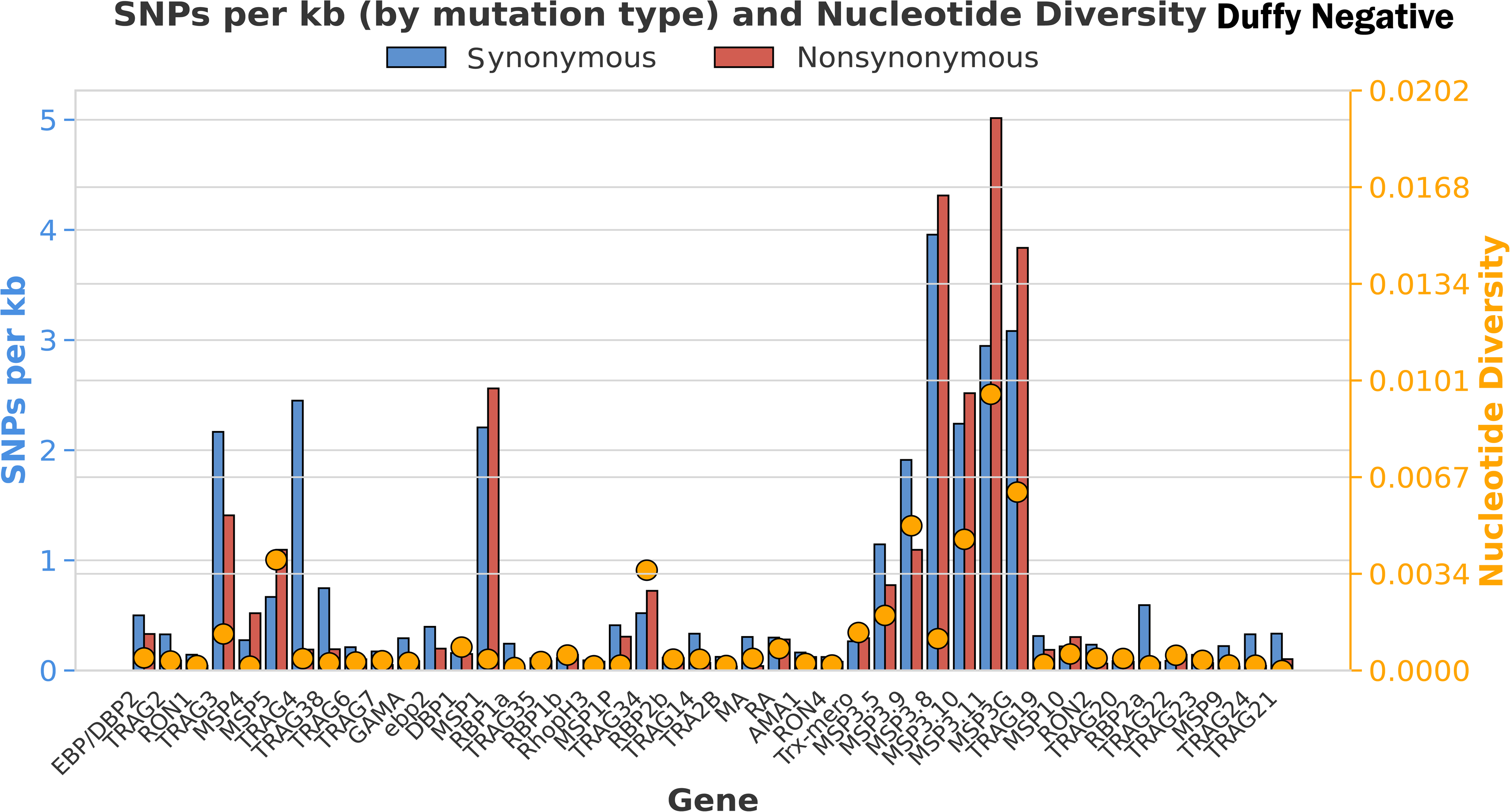
Distribution of synonymous and nonsynonymous SNPS across all 14 chromosomes of the nuclear genome. Consistently higher proportions of synonymous SNPs to nonsynonymous SNPs with more mutations were detected in Duffy positive than Duffy negative samples. Chromosomes 9, 10, and 12 have the higher portion of mutations across all chromosomes in both Duffy groups, with chromosomes 3 and 6 the least polymorphic.

For the 43 erythrocyte-binding genes, 13,527 SNPs (8,018 synonymous and 5,509 nonsynonymous) were detected in Duffy-positive samples, with an average nucleotide diversity (π) of 4·39×10^−3^ (Supplementary Table 2). The least polymorphic loci, including *PvRBP1a*, *PvTRA*2B, *PvMA*, and *PvRON*1, accounted for only 0·3-0·5% of total SNPs and exhibited fewer than 1 SNP per kb. By contrast, markedly fewer SNPs (6,212 total; 3,168 synonymous and 2,877 nonsynonymous) were detected in Duffy-negative samples, with lower average nucleotide diversity (π=0·98×10^−3^; Supplementary Table 2). *PvRBP1a*, *PvMSP9*, *PvTRA2b* and *PvMA* were also shown to be conserved in Duffy-negative samples (accounted for 0·2-0·4% of all SNPs), suggesting broad functional constraint. In both Duffy-negative and Duffy-positive *Pv*, the highest polymorphisms were observed in *PvTRAg3 and PvTRAg4,* as well as members of the *PvMSP3* multigene family (*PvMSP*3G, *PvMSP*3.11, *PvMSP*3.10, and *PvMSP*3.8; Figure 2). These loci accounted for 46% of all SNPs in Duffy-positive and 43% in Duffy-negative *Pv*.

**Figure 2.**
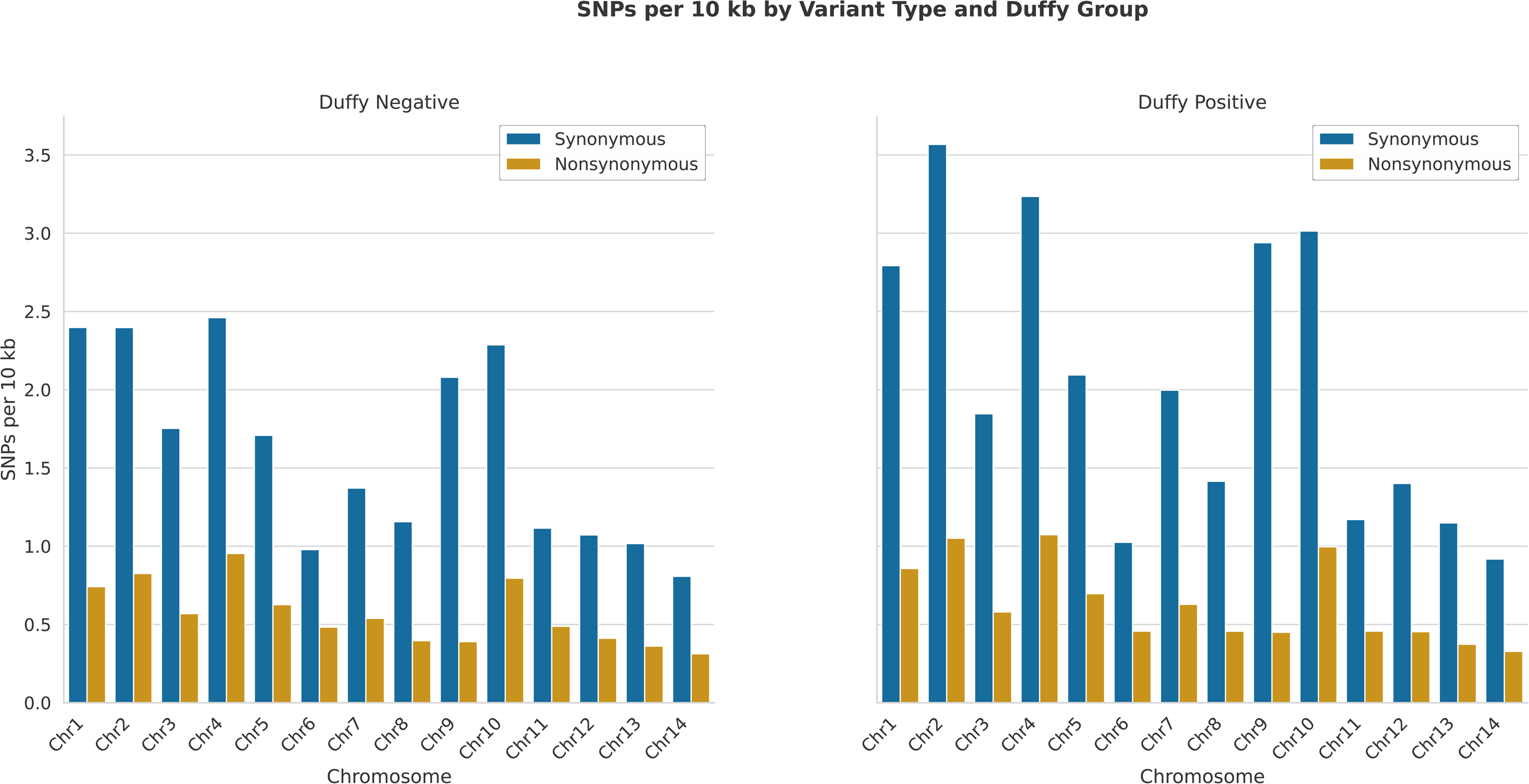
Distribution of synonymous and nonsynonymous SNPs in 43 erythrocyte binding genes, as well as nucleotide diversity in each gene. Genes in the *PvMSP*3 multigene family had higher levels of SNPs and nucleotide diversity relative to other erythrocyte binding genes. *PvMSP*3G, *PvMSP*3.11, and *PvMSP*3.10 showed the highest levels of polymorphisms in both Duffy groups.

### Differential mutation profiles of key invasion ligands

To investigate fine-scale divergence in key invasion ligands, we compared nonsynonymous codon substitutions between Duffy-positive and Duffy-negative *Pv* isolates (Figure 3). In *PvDBP*1, 35 codon substitutions were identified in Duffy-positive *Pv*, with individual mutation frequencies ranging from 12-51%. Fifteen of these substitutions were located within the binding domain of region II. Most notably, over 50% of Duffy-positive samples had mutations at codons G175E, S263R, K402S, and I458K, all within the binding domain (Figure 3A). In contrast, fewer than 5% of Duffy-negative samples had these same mutations, and only 20 distinct codon substitutions were detected in 5-10% of the isolates. In *PvEBP/DBP*2, 14 codon substitutions were identified in Duffy-positive *Pv,* with frequencies ranging from 10-43% (Figure 3B). Eight of these occurred in the binding domain, where nearly half of samples exhibited mutations at codons K306E and I421T. Among Duffy-negative samples, only four substitutions were detected, each in 5-7% of the samples. The same variants K306E and I421T were found in <5% of the samples.

**Figure 3.**
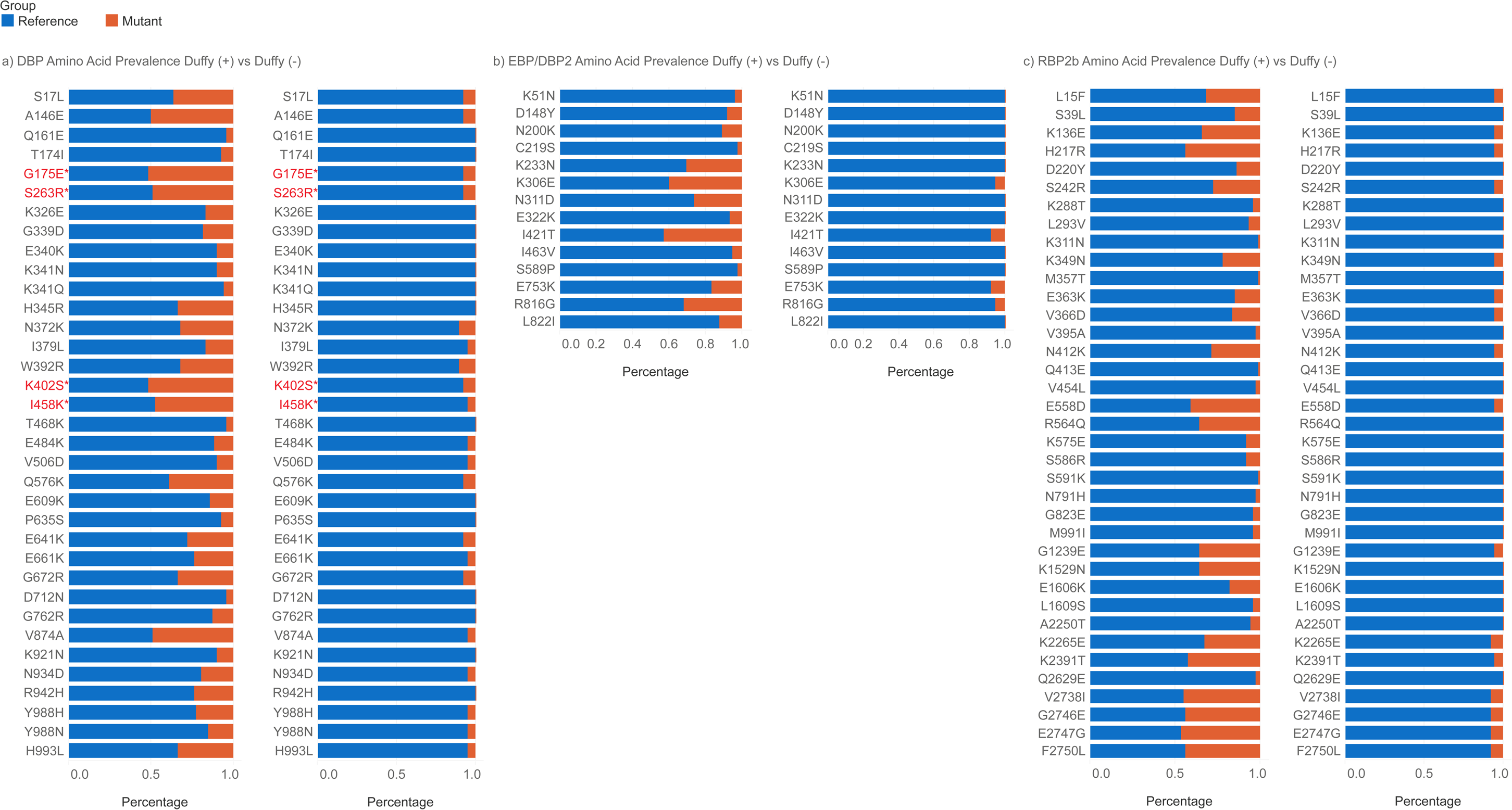
Amino acid variations across three erythrocyte binding genes including *PvDBP*1, *PvEBP/DBP*2, and *PvRBP*2b, all of which are known to involve in *Pv* invasion of Duffy-positive erythrocytes. Asterisks denote codons located in the binding regions. In *PvDBP*1, over 50% of Duffy positive samples were observed with mutation at four codon positions including G175E, S263R, K402S, and I458K (in red), all of which were in the binding region. In contrast, less than 10% of Duffy negative samples had these mutations at the same position. No mutations were detected in binding region of *PvEBP/DBP*2 nor *PvRBP*2b that were markedly different between the Duffy-positive and Duffy-negative samples.

A similar pattern was observed in *PvRBP*2b, of which 37 codon substitutions were identified in Duffy-positive *Pv,* with frequencies ranging from 1-46% of the isolates; 23 of which occurred in the binding domain (Figure 3C). 217R and 558D were two most prevalent substitutions detected in up to 50% of the Duffy-positive *Pv*. In contrast, only 16 substitutions were detected in 5-8% of Duffy-negative *Pv*, including eight mutations within the binding region that found in <5% of the samples. No mutations were detected in binding region of *PvEBP/DBP*2 nor *PvRBP*2b that were unique in the Duffy-negative isolates, indicating an overall genomic conservation of these invasion ligands in this group. For the other erythrocyte-genes *PvMSP1*, *PvRBP1a*, and *PvRBP2a*, four codon substitutions, including E1692K (*PvMSP1*), P552Q (*PvRBP1a*), E438G, and M511K (*PvRBP2a*), were detected in >50% of the Duffy-positive but <10% of Duffy-negative isolates (Supplementary Figure 1A-E). These mutations were located at the binding domains of the respective ligand. Collectively, *PvDBP*1, *PvEBP/DBP2*, and *PvRBP*2b were genetically more conserved in Duffy-negative than Duffy-positive *Pv*, with *PvEBP/DBP*2 being the most conserved gene, implicating limited diversification under strong functional constraint.

### Distinct signatures of positive selection in Duffy-positive and Duffy-negative *Pv*

To identify genomic regions under recent positive selection, we calculated the integrated haplotype score (iHS) across the 14 *Pv* chromosomes with respect to each Duffy group. Duffy-positive *Pv* showed significant iHS signals in six distinct gene regions (*p*<0·001; Figure 4A). Among these, two loci including chloroquine resistance-associated protein 1 (*Cg*1; PvP01_0109400) and 26S proteasome regulatory subunit (*RPN*2; PVP01_1456300) exhibited the strongest signals of recent positive selection. The region encompassing *Cg*1 has been linked to the chloroquine resistance locus in *Pf*, although chloroquine resistance has not been experimentally confirmed in *Pv*. Similarly, variation in *RPN*2 has been linked to altered artemisinin response in *Pf*, but the association remains tentative and not confirmed outside *Pf*, where *Kelch*13 is the principal determinant of artemisinin resistance (37). ^55^

**Figure 4.**
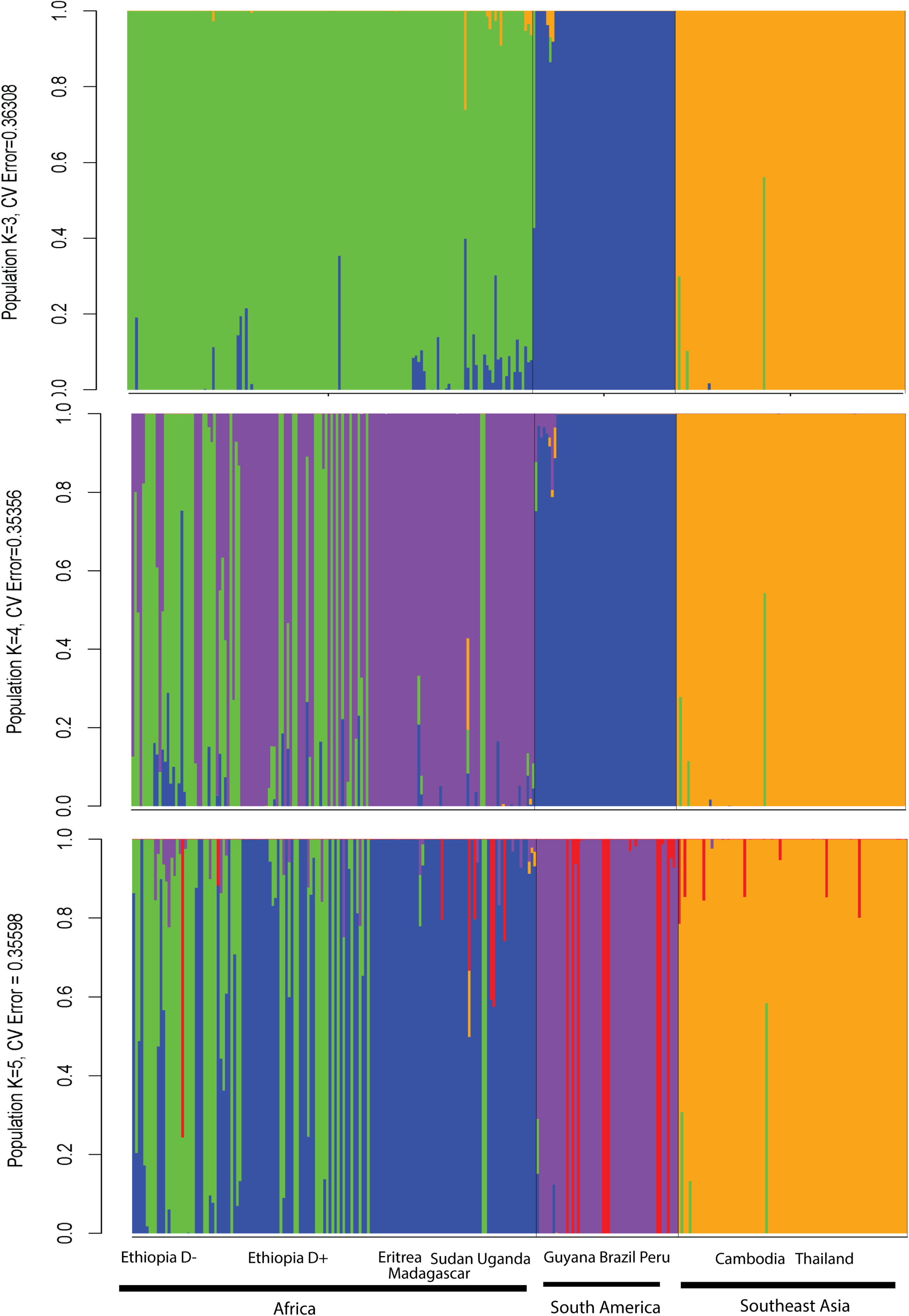
Bar plots based on ADMIXTURE analyses of SNP profiles show clustering patterns of sample data collected from Africa, Southeast Asia and South America. results for *K*=3, 4 and 5 coupled with their respective cross validation errors. Three genetic clusters were consistently observed for each *K*, one for each continental region. At *K*=5, high levels of admixture was seen among the African and South American *Pv*. No clear differentiation was detected between Duffy-positive and Duffy-negative samples.

In *Pv* from Duffy-positive samples, 72% had the *PvCg*1mutant allele (G to A) and 69% had the *PvRPN*2 mutants, each supported by sequencing depth greater than 10x. In addition, PVP01_0533300 (*PvSTP1*) on chromosome five, a gene structurally related to the surface-associated interspersed protein (*surf*) family and expressed on the surface of infected erythrocytes and merozoites, showed strong selection signals (*p*<0·001) (38).^38^ Most of the SNP variants on chromosome 9 that show signals of positive selection occurred in intergenic regions flanking hypothetical proteins, with the exception of the membrane-associated erythrocyte binding-like protein (*MAEBL*; PVP01_0948400), an invasion ligand implicated in merozoite attachment and red blood cell entry (39, 40). In contrast, Duffy-negative *Pv* displayed fewer genomic locations under significant positive selection (Figure 4B). Significant signals were found in an unknown conserved protein (PvP01_1322000, *p*<0·001) and a plastid replication-repair enzyme (PVP01_1337500, *p*<0·001) on chromosome 13, and the AP-5 complex subunit β1 (PVP01_1438600, *p*<0.001) on chromosome 14.

### Population structure and admixture analyses reveal limited differentiation

The most probable number of genetic clusters identified by *K*-fold cross validation scores in the admixture analysis were *K*=3 (error=0.363), *K*=4 (error=0.354), and *K*=5 (error=0.356), indicating that three to five ancestral components best explained global *Pv* population structure (Figure 5). At *K*=3, *Pv* in Duffy-positive and Duffy-negative Ethiopians shared the same genetic cluster that included *Pv* from Sudan and Uganda, suggesting a high degree of genetic similarity across East African populations. Most Southeast Asian *Pv* formed a distinct clustered, though a small subset showed admixture with the African lineages. In contrast, the South American *Pv* were genetically distinct from both the African and Southeast Asian isolates, reflecting deep continental divergence. Similar clustering patterns were observed at *K*=4 and *K*=5, where samples were segregated into distinct geographical clusters corresponding to Southeast Asia, South America, and two African subgroups (Figure 5), consistent with previous studies (24). Within Africa, one cluster encompassed isolates from Eritrea, Uganda, Sudan, and Madagascar, while a second cluster encompassed isolates from Ethiopia, some of which were genetically distinct from their regional counterparts. Despite this sub-structuring pattern, *Pv* from Duffy-negative Ethiopians did not form a distinct genetic cluster from those in Duffy-positives (Figure 5), suggesting a shared evolutionary origin within East Africa.

**Figure 5.**
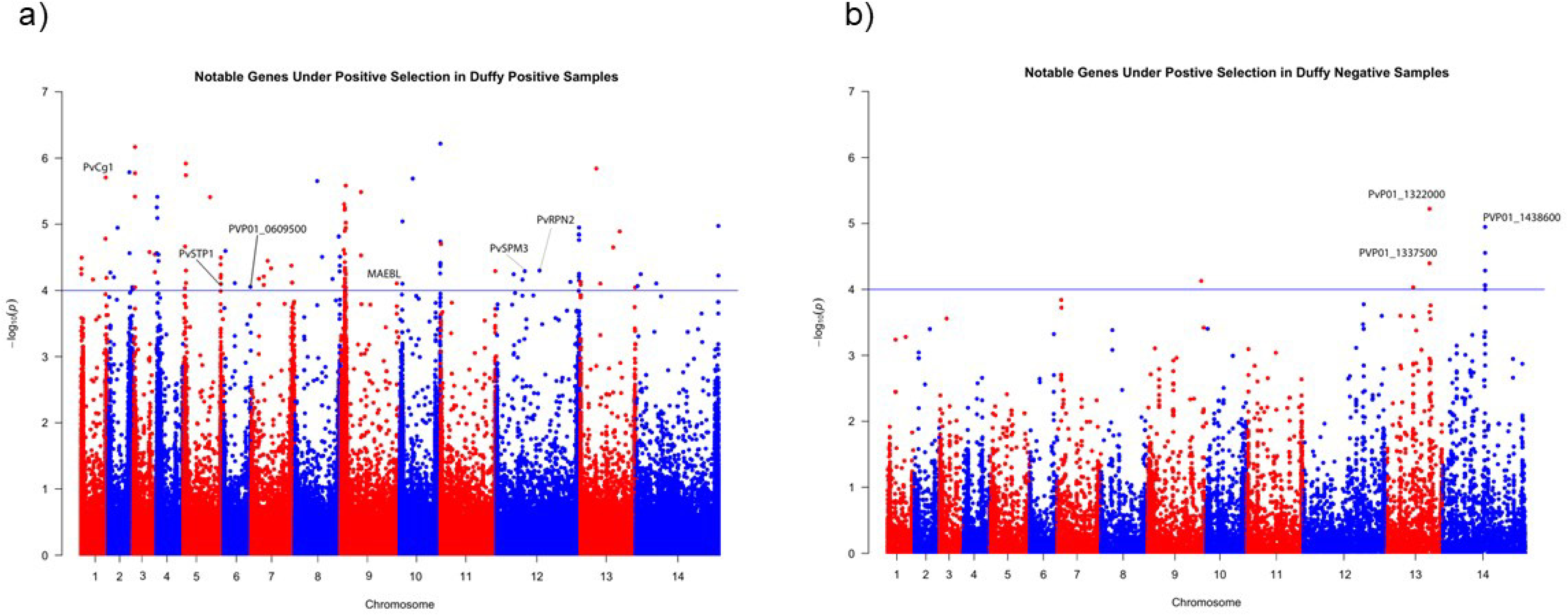
Signals of positive selection across the 14 chromosomes in both Duffy positive and negative Ethiopians. Genes that showed significant signals of positive selection in Duffy positive samples included MAEBL, Cg1, STP1, SPM3 and RPN2, with genes Cg1 and RPN2 involved with drug resistant and MAEBL being a potential erythrocyte binding gene. Genes with significant positive selection in Duffy negative samples include a plastid replication repair enzyme and the AP-5 complex subunit beta one.

## Discussion

This study reveals distinct yet interconnected patterns of genetic variation, selection, and population structure of *Pv* in Duffy-positive and Duffy-negative populations in Ethiopia, providing new insights into parasite adaptation in Africa. We identified a substantial number of high-quality SNPs, with more than twofold higher SNP count in Duffy-positive *Pv*, consistent with a larger and more heterogeneous parasite reservoir in Duffy-positive population. Similar patterns of genetic diversity have been observed in other East African *Pv* based on microsatellites, supporting the presence of genetically diverse lineages in this region (41, 42). The pronounced polymorphisms in Duffy-positive parasites, especially on chromosomes 9, 10, and 12, were largely driven by variability within the *PvMSP3* multigene family, which encodes merozoite surface proteins implicated in immune evasion and erythrocyte invasion (43). Such polymorphisms may reflect an evolutionary arms race between host immunity and parasite adaptability, where recombination and immune selection act as major forces shaping local *Pv* populations. In contrast, Duffy-negative individuals account for only one-third of the general population in Ethiopia and other East African countries (4). The smaller host populations may impose bottlenecks, limiting recombination, reducing effective population size, and resulting in lower genomic diversity observed in Duffy-negative *Pv*. Despite this difference, the overall population clustering patterns suggest that both Duffy groups share recent common ancestry and ongoing gene flow, reflecting a unified parasite gene pool circulating within East Africa.

Among the 43 erythrocyte-binding genes examined, *PvDBP1*, *PvEBP/DBP2*, *PvRBP2b,* and some members of the *PvTRAg* family were among the most conserved in both Duffy groups. These genes encode key ligands for erythrocyte recognition, for instances, *PvEBP/DBP2* region II binds to the LHR-A subdomain of CR1 while *PvRBP2b* (residues 161-1454) engages the transferrin receptor 1 (TfR1 or CD71) on reticulocytes (11, 16, 17, 44), although recent studies indicated that binding inhibitors blocking the TfR1 receptor showed no inhibition of *Pv* invasion in the Cambodian isolates (45). The functional significance of the PvRBP2b/TfR1 pathway may be varied by parasite strains. Conservation of these genes in Duffy-negative *Pv* suggests their essential role in maintaining parasite functions for successful invasion of erythrocytes. It has been hypothesized that unique mutations in *PvDBP*1 may offer the parasites invasion capability to Duffy-negative erythrocytes. While our earlier study revealed a few unique *PvDBP*1 mutations in two Duffy-negative *Pv* samples (10), the expanded dataset in this study revealed no Duffy group-specific variants, reinforcing the likelihood that PvDBP1*-*DARC interaction remains functionally relevant in Duffy-negative hosts. This notion aligns with evidence that DARC is transiently expressed on immature erythroid precursor cells within bone marrow and splenic niches of Duffy-negative individuals (46), suggesting that invasion may occur in early-stage reticulocytes before their release into circulation (46, 47).

Genes in the *RBP* and *TRAg* (with the exception of *TRAg*34) multigene families are generally conserved across both Duffy groups compared to the *MSP*, potentially reflecting immune selection. *Pv*-infected individuals from Northern India have shown strong humoral immune responses against conserved regions of *PvTRAg* (48, 49). *PvTRAg*34, one of four *PvTRAg* genes that bind to erythrocytes and shares the same erythrocyte receptors with *PvTRAg*33.5, (50–52) displayed elevated polymorphism potentially driven by balancing selection linked to host immunity. Recent transcriptomic analyses suggested that *Pv* from different geographical regions may utilize different invasion repertoires (53). For example, *PvDBP1* and *PvEBP/DBP2* were highly expressed in the Cambodian but less so in the Ethiopian and Brazilian *Pv*, whereas *PvRBP2a* and *PvRBP2b* exhibited higher expression in the Ethiopian and Cambodian isolates. These expression profiles, together with the conserved nature of *PvDBP* and *PvTRAg* gene families, suggest that parasite invasion strategies could be shaped by regional host genetics and transmission intensity. The reduced diversity of invasion ligands in Duffy-negative isolates may imply stronger purifying selection and/or constrained host-receptor availability, maintaining essential invasion functions under reduced genetic flexibility.

Our selection analyses further demonstrated that adaptive pressures differ markedly between Duffy-positive and Duffy-negative *Pv*. In Duffy-negative *Pv*, significant positive selection were detected in *PvCG*1 and *PvRPN*2, genes link to chloroquine and artemisinin resistance in *Pf* (37, 54), as well as in *PvMAEBL*, an invasion ligand implicated in sporozoite attachment to the salivary glands of *Anopheles* mosquitoes (55, 56). In contrast, Duffy-negative *Pv* exhibited significant selection in genes encoding a plastid replication-repair enzyme and the AP-5 complex subunit β1. AP-5 encodes a component of the adaptor protein complex involved in intracellular membrane trafficking and formation of post-Golgi vesicles (57–60), suggesting possible adaptations in intracellular trafficking or organellar maintenance in Duffy-negative *Pv* (61). These adaptations may facilitate parasite survival in altered intracellular environments and ensure efficient trafficking of proteins and nutrients essential for erythrocyte invasion under restricted receptor conditions (57, 62). Collectively, these findings indicate that while Duffy-positive *Pv* shows adaptive signals in drug-response and erythrocyte-invasion genes, Duffy-negative parasites exhibit selective pressures in genes associated with cellular transport and organellar homeostasis. This contrast implies differential evolutionary trajectories reflecting host-dependent ecological and molecular constraints.

Despite the difference in the level of genomic diversity, *Pv* in Duffy-positive and Duffy-negative individuals were not genetically distinct but clustered together with other East African *Pv* lineages from Sudan and Uganda, consistent with previous findings based on microsatellites (41) and suggests a shared ancestry and/or recent gene flow between Duffy populations and across regions (23, 26). No significant difference was observed between Duffy positive and Duffy negative *Pv*, Nevertheless, these African isolates remain distinct from the Southeast Asian and South American lineages, aligns with prior global studies showing three distinct ancestral clusters, respectively, in Asia, Africa, and South America (24). A recent microhaplotype-based analysis showed markedly reduced genetic diversity in *Pv* from the Horn of Africa compared with other geographical regions (63). Consistent with the parasite evolutionary origin in Asia (24, 63), diversity was the highest across Asian *Pv* populations and declined progressively with increasing geographical distance from Asia. Reduced diversity and elevated linkage disequilibrium in the African isolates further suggest a serial founder effect following *Pv* migration from Asia, with subsequent local adaptation to Duffy-negative hosts (4, 41, 63).

While our study offers valuable insights into the genomic features of Duffy-negative *Pv*, there were few limitations. First, Duffy-negative individuals from Ethiopia alone may limit the generalizability of our findings. Expanded samples from multiple African countries would provide a more comprehensive view of genetic variations and adaptive strategies of *Pv* in diverse host populations. Second, our analyses focused primarily on SNP-level variations, without exploring other types of genomic diversity such as copy number changes, structural variants, and gene expression differences. This may undermine important mechanisms underlying *Pv* adaptation in Duffy-negative hosts. Third, while we detected signals of recent positive selection in Duffy-negative Ethiopians, the functional implications of these candidate loci remain unclear. Experimental validation through *in vitro* studies or model systems would offer better understand of their roles in infection, invasion, or immune evasion. Finally, this study examined solely on parasite genomics, but host genetic factors that could significantly influence infection dynamics and parasite adaptation were not evaluated. Future research integrating both host and parasite genomic data would provide a more holistic understanding of the biological interactions shaping *Pv* evolution in Africa.

## Conclusion and Future Directions

Our study provides the first genome-wide comparison of *Pv* diversity in Duffy-negative and Duffy-positive individuals, uncovering signatures of adaptation and conservation that underpin parasite persistence in Africa. The remarkably conserved genomes in Duffy-negative *Pv*, together with positive selection in genes linked to organellar function and vesicular trafficking, point to functional adaptation that may optimize parasite survival in hosts previously considered to be less susceptible. The lack of genetic structure between the two Duffy groups implies frequent gene flow and a shared evolutionary origin among Ethiopia *Pv* lineages. Further studies expanding sampling across Africa and integrating parasite and host genomic data will be essential to unravel the molecular mechanisms of Duffy-independent invasion. Functional validation of candidate genes identified here, particularly *PvEBP/DBP2*, *PvRBP2b*, and the AP-5 complex, will illuminate the cellular pathways exploited by *Pv* to invade Duffy-negative erythrocytes. Such insights are critical for refining malaria surveillance, developing next-generation diagnostics, and guiding elimination strategies tailored to the changing epidemiology of *Pv* in Africa.

## Acknowledgements

We thank the field team from Jimma University for their technical assistance; the communities and hospitals for their support and willingness to participate in this research; Drs. Joshua Mell and Azad Ahmed from the Drexel Genomics Core Facility and SKCC Share Genomics Resources for consultation and sequencing services; and undergraduate students at Drexel University for assistance with the experiments.

---

**Supplementary Figures 1.** Amino acid variations of erythrocyte binding gene candidates including (A) *PvMSP*1, (B) *PvRBP*1a, (C) *PvRBP*1b, (D) *PvRBP*2a, and (E) *PvTRAg38.* Asterisks denote codons located in the binding regions. Codons in red indicate mutations in the binding region that were found in over 50% of Duffy positive samples but less than 10% of Duffy negative samples at the same position. They included E1692K in *PvMSP1*, P552Q in *PvRBP1a*, and E438G and M511K in *PvRBP2a*.

