## Supplementary figures and images for "Adaptive evolution of *Plasmodium vivax* in Duffy-negative hosts: insights from East African genomes"

### Supplementary Figures

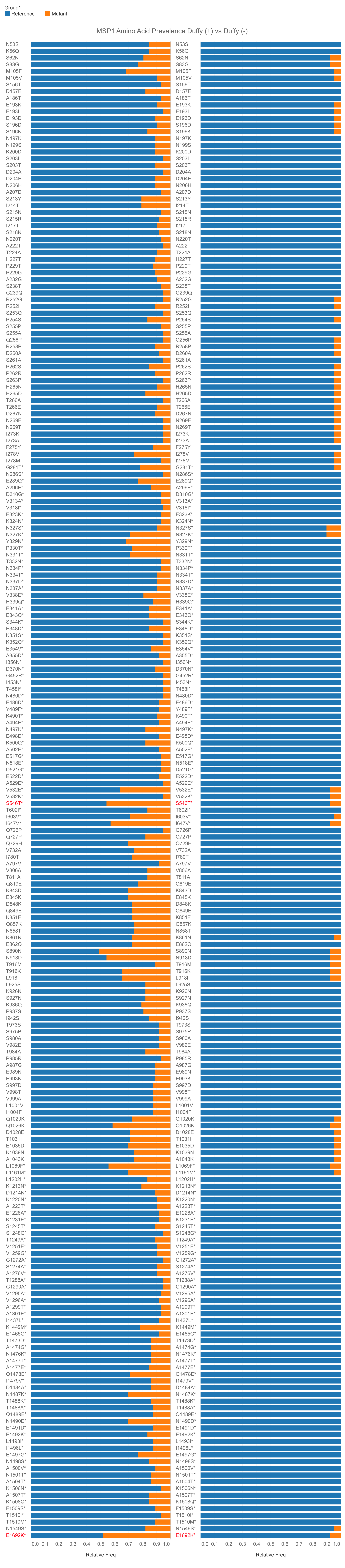

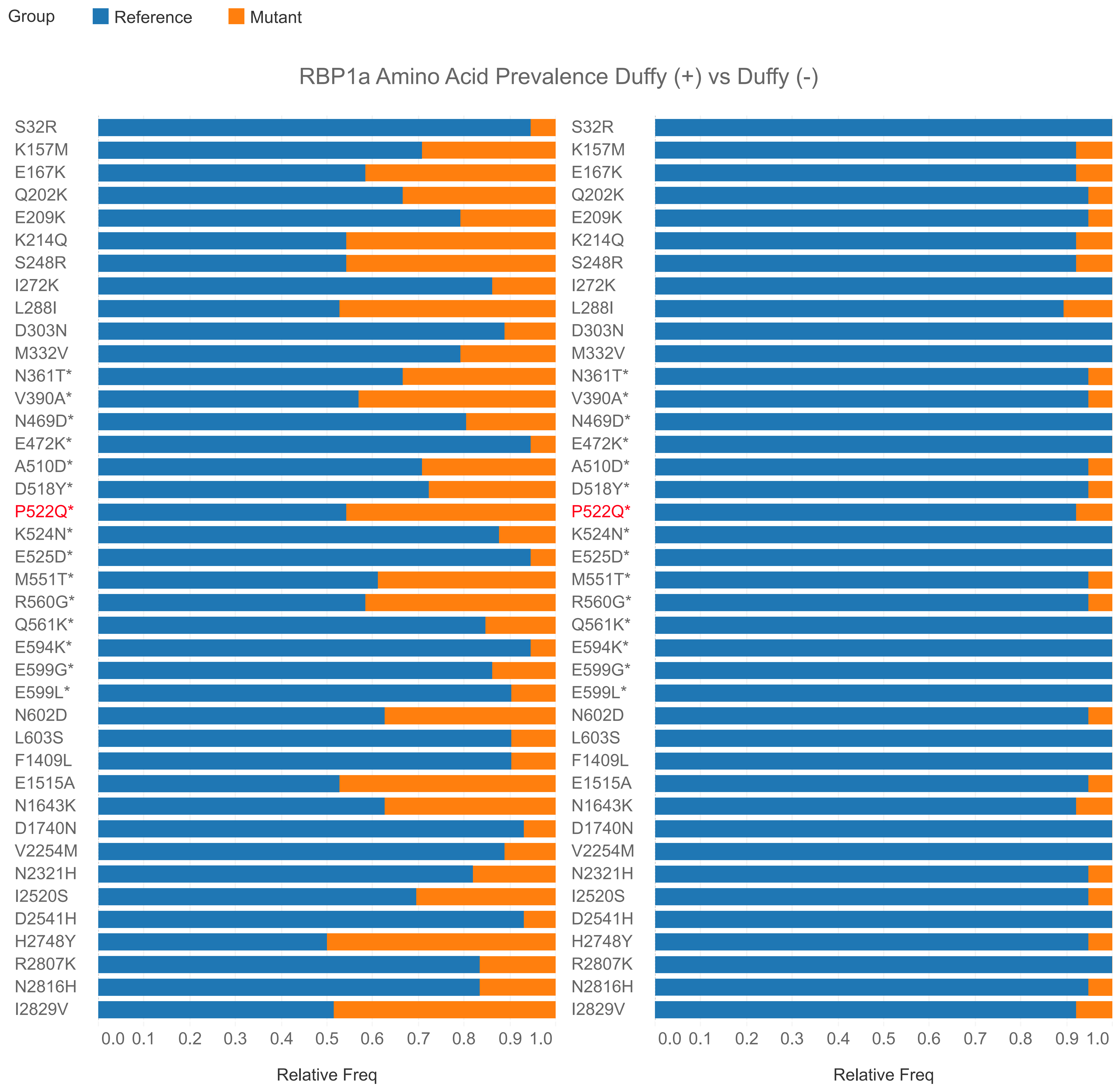

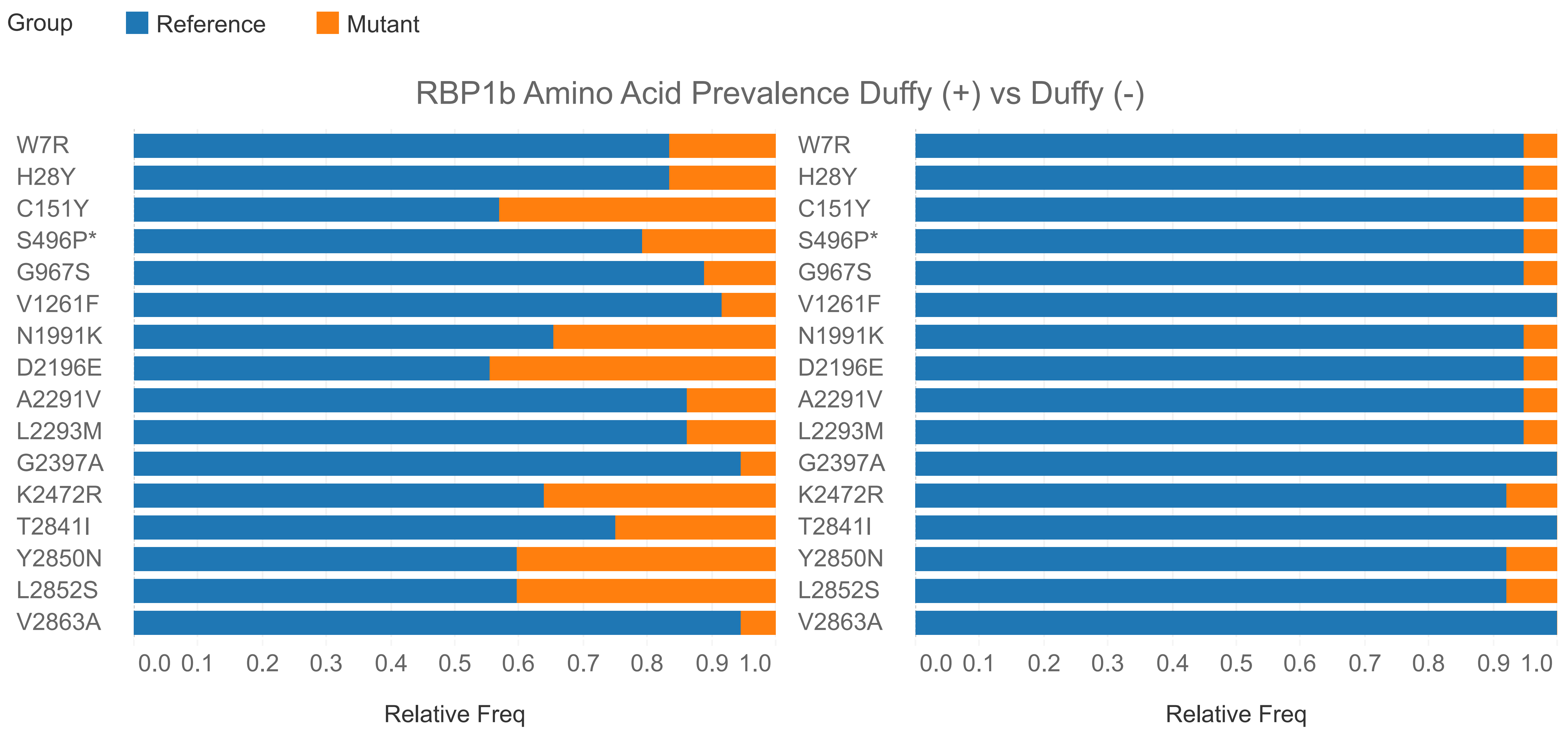

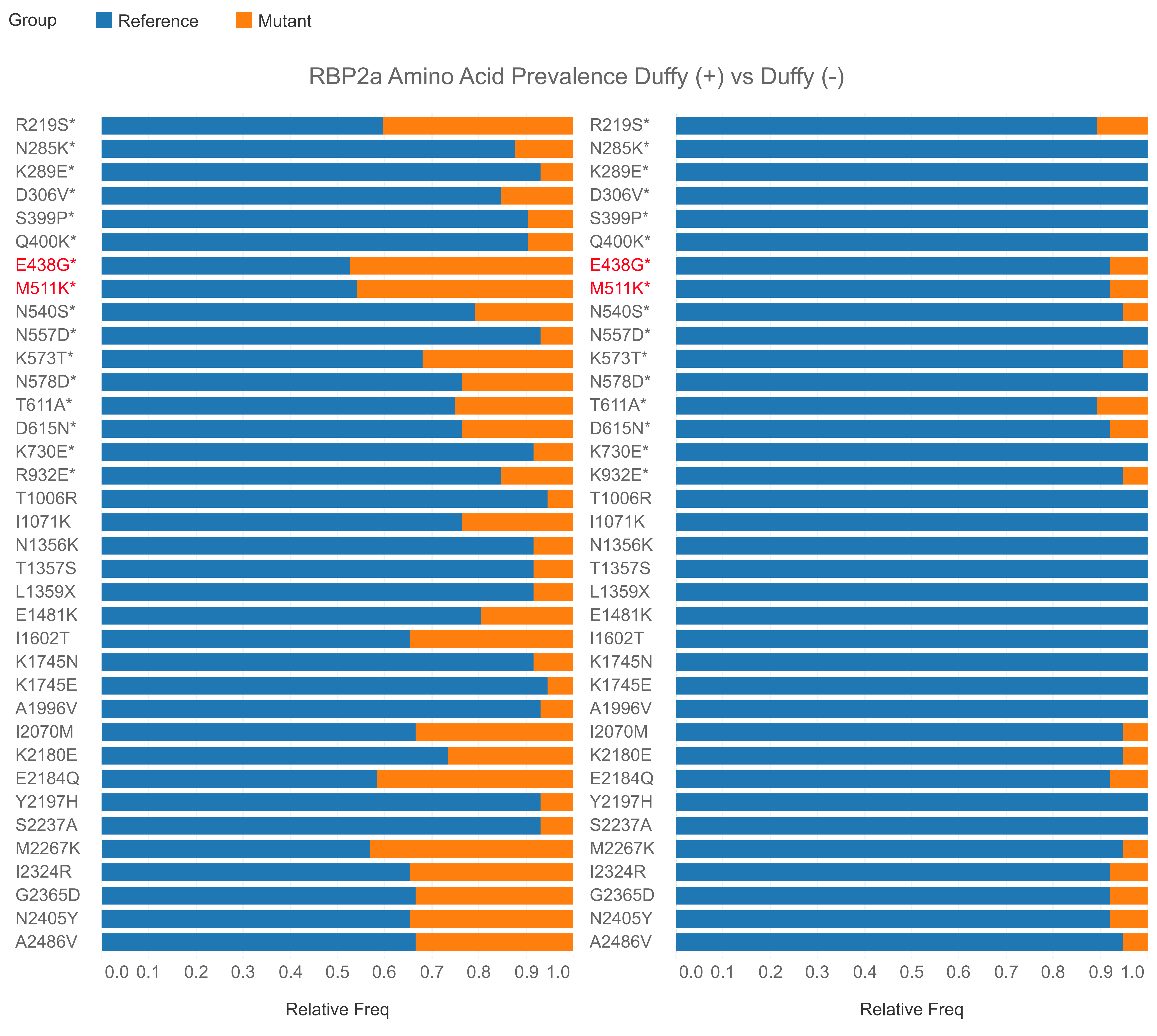

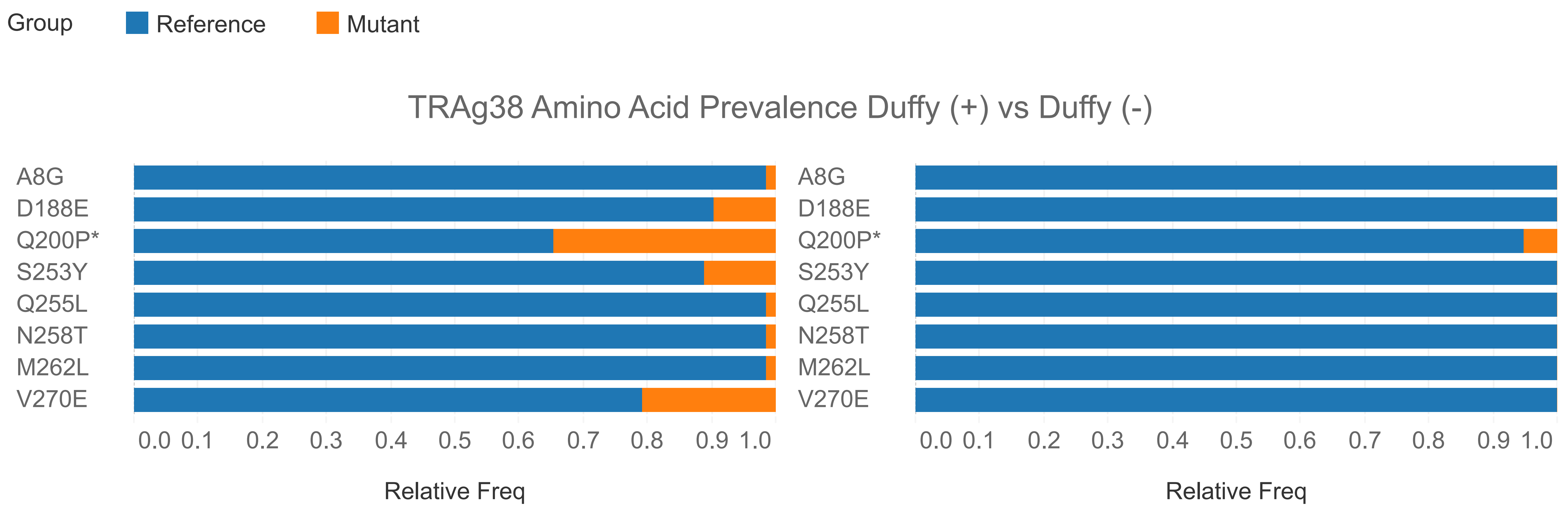
